# Genetic Variation, Not Cell Type of Origin, Underlies Regulatory Differences in iPSCs

**DOI:** 10.1101/013888

**Authors:** Courtney L. Kagan, Nicholas E Banovich, Bryan J Pavlovic, Kristen Patterson, Irene Gallego Romero, Jonathan K Pritchard, Yoav Gilad

## Abstract

The advent of induced pluripotent stem cells (iPSCs)^1^ revolutionized Human Genetics by allowing us to generate pluripotent cells from easily accessible somatic tissues. This technology can have immense implications for regenerative medicine, but iPSCs also represent a paradigm shift in the study of complex human phenotypes, including gene regulation and disease^2-5^. Yet, an unresolved caveat of the iPSC model system is the extent to which reprogrammed iPSCs retain residual phenotypes from their precursor somatic cells. To directly address this issue, we used an effective study design to compare regulatory phenotypes between iPSCs derived from two types of commonly used somatic precursor cells. We show that the cell type of origin only minimally affects gene expression levels and DNA methylation in iPSCs. Instead, genetic variation is the main driver of regulatory differences between iPSCs of different donors.

---

Research on human subjects is limited by the availability of samples. Practical and ethical considerations dictate that functional molecular studies in humans can generally only make use of frozen post mortem tissues, a small collection of available cell lines, or easily accessible primary cell types (such as blood or skin cells). The discovery that human somatic cells can be reprogrammed into a pluripotent state^6,7^ and can then be differentiated^8^ into multiple somatic lineages, has the potential to profoundly change human research by providing us access to a wide range of cell types from practically any donor individual.

Though much progress has been made since the initial development of iPSC reprogramming technology, and human iPSCs have been used in a wide range of studies, the usefulness of iPSCs as a model system for the study of human phenotypes is still extensively debated^9-11^. The principal issue is the extent to which reprogrammed iPSCs retain epigenetic and gene expression signatures of their cell type of origin. A residual epigenetic signature of the original precursor cell in the reprogrammed iPSCs is often referred to as ‘epigenetic memory’^12^.

The common view, established by a few early studies in mice and humans, is that epigenetic memory is a significant problem in iPSCs^10, 12-18^ In mice, methylation profiles in iPSCs and in the precursor somatic cells from which the iPSCs were generated were found to be more similar than expected by chance alone^12, 14^ The extent of this similarity, however, could not be benchmarked against genetic diversity because the somatic cells and the iPSCs were all from genetically identical mice. In turn, in humans, methylation profiles in iPSCs reprogrammed from different somatic cell types were found to be quite distinct from each other^15, 16^ However, the somatic cells were provided by different donor individuals, hence epigenetic memory and differences due to genetic diversity were confounded.

An additional aspect to consider is that the notion that epigenetic memory in iPSCs may be an important concern was established by studies that considered iPSCs generated using retroviral vectors^12, 14-16^. Retroviral reprogramming of iPSCs leads to random integrations that vary in copy number and genomic location across lines. The process of random integration could have appreciable effects on gene regulation, especially if the location and copy number of the integrations are cell type specific (e.g., due to differences in chromatin accessibility across cell types). In contrast to retroviral reprograming, the more recent episomal approaches to establish iPSCs are associated with much lower rates of genomic integration^19, 20^

Indeed, one recent study has concluded that when properly controlling for genetic variation and using integration free methodology to establish iPSCs, the effect of cell type of origin on gene expression is low compared to inter-individual genetic contributions^21^. However, this study did not consider matched epigenetic markers, the supposed drivers of the suspected phenomenon of residual cell type of origin memory in reprogrammed iPSCs.

We thus designed a study to directly and effectively address this issue. We focused on two cell types that are the source for the majority of human iPSCs to date, and the most easily collected tissue samples from humans: skin fibroblasts, and blood cells. Specifically, we collected skin biopsies and blood samples from four healthy Caucasian individuals (two males and two females). Dermal fibroblasts were isolated from dissociated skin biopsies and maintained in culture until reprogramming. We isolated the buffy coat from whole blood and subsequently used Epstein–Barr virus to transform B cells into immortalized lymphoblastoid cell lines (LCLs), a common cell type used in genomic studies.

We used an episomal reprogramming approach^19^ to independently generate iPSCs from the LCLs and fibroblasts of each individual, three replicates from the LCLs (to study technical variance) and one from the fibroblasts (to study epigenetic memory; Fig. 1A). We employed a wide range of quality control analyses and functional assays to demonstrate that all iPSCs were fully pluripotent, that they expressed endogenous, but not exogenous, pluripotency factors, that the iPSCs were free of vector integrations, and that iPSCs established from LCLs did not retained traces of integrated EBV (see supplementary methods; Supplementary Figs. 1-4).

**Figure 1.**
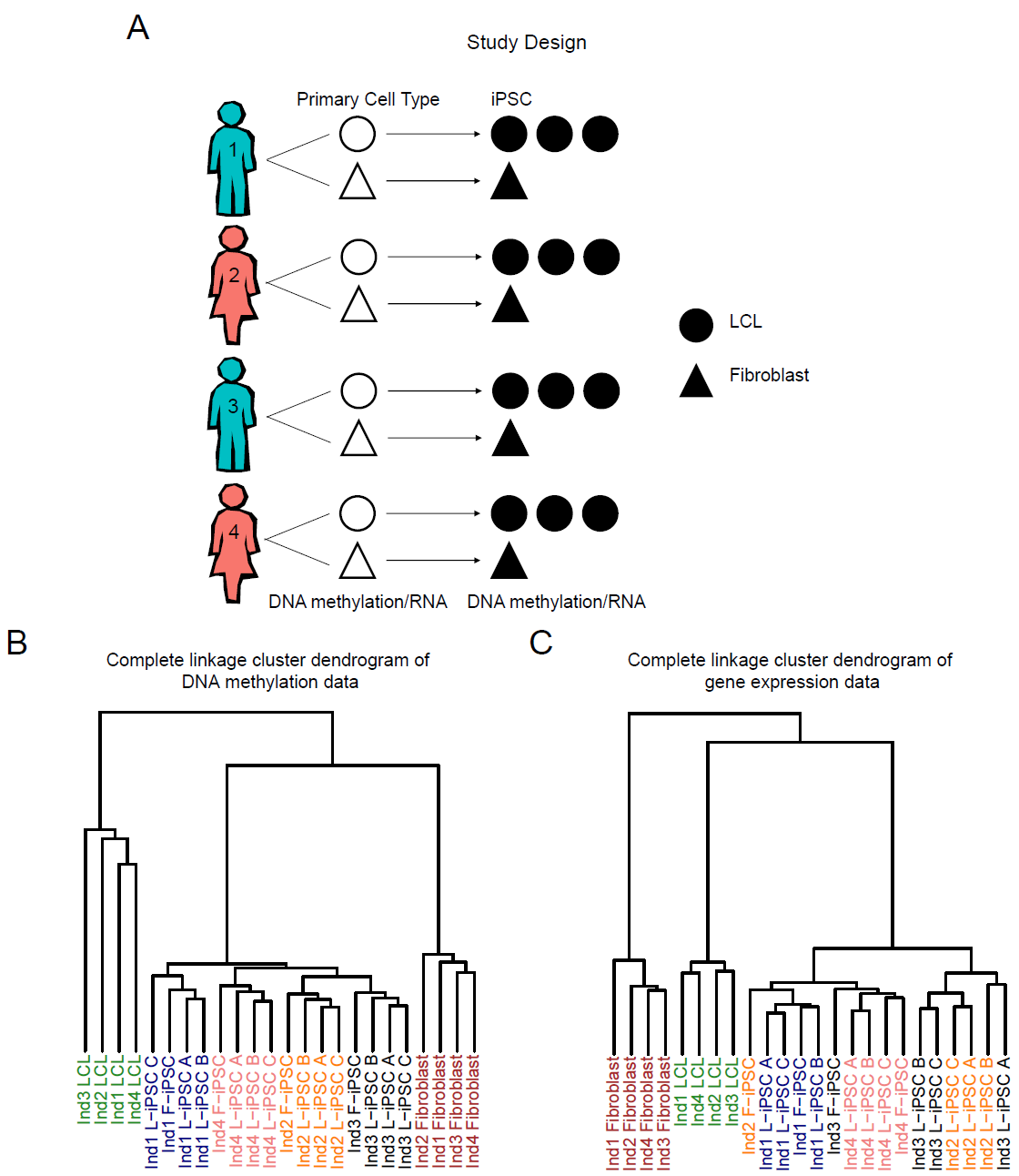
Study design and clustering of samples based on DNA methylation or gene expression data. (a) Study design schematic. Three independent iPSC lines were generated from LCLs and one from fibroblasts. Hierarchical clustering using the complete linkage method and Euclidean distance from autosomal loci for (b) DNA methylation data (n = 445,277 probes) and (c) gene expression data (n = 10,648 autosomal genes).

Once the quality of the iPSCs was confirmed, we extracted RNA and DNA from LCLs, fibroblasts, LCL derived iPSCs (L-iPSCs), and fibroblast derived iPSCs (F-iPSCs) from all four individuals (Supplementary Table 1). We then used the Illumina Infinium HumanMethylation450 array and the Illumina HumanHT12v4 array to measure DNA methylation and gene expression levels, respectively. Our data processing approach is described in detail in the supplementary methods. Briefly, considering the methylation data, we first excluded data from loci that were not detected either as methylated or unmethylated (no signal; detection *P* > 0.01) in more than 25% of samples. We then applied a standard background correction^22^ and normalized the methylation data using SWAN^23^ (Supplementary Fig. 5), which accounts for the two different probe types in the platform. Finally, we performed quantile normalization (Supplementary Fig. 6A/B). Following these steps we retained methylation data from 455,910 CpGs. Considering the expression data, we first excluded probes whose genomic mapping coordinates overlapped a known common SNP. We then retained all genes that were detected as expressed in any cell type in at least three individuals (Supplementary Fig. 7). We then quantile normalized the gene expression data (Supplementary Fig. 6C/D). Following these steps we retained expression data for 11,054 genes.

To examine overall patterns in the data, we initially performed unsupervised clustering based on Euclidean distance. As expected, using gene expression or methylation data, samples clustered based on cell type (LCLs, fibroblasts, and iPSCs) without exception. Interestingly, using the methylation data, iPSCs clustered perfectly by individual, not cell type of origin (Fig. 1B). This pattern was largely recapitulated when we considered the gene expression data (Fig. 1C), however the clustering was not as robust (most likely because there are far fewer expression measurements than methylation measurements). Given the large number of sites interrogated (particularly on the methylation array), we also examined the clustering of iPSCs using only the top 1,000 most variable measurements across lines, similar to the approach of Kim et al. 2011^16^ Our clustering remained largely unchanged using this subset of variable sites for both methylation data (Supplementary Fig. 8A) and expression data (Supplementary Fig. 8B). Clustering based on pairwise Pearson correlations rather than Euclidian distance produced identical results (Supplementary Fig. 8C/D).

We next considered methylation and expression patterns at individual loci and genes, respectively. We first focused on differences in CpG methylation between the cell types. Using limma^24^ (see supplementary methods), we identified 190,356 differentially methylated (DM) CpG loci between LCLS and fibroblasts (FDR of 5%). Similarly, we identified 310,660 DM CpGs between LCLs and L-iPSCs and 226,199 DM loci between fibroblasts and F-iPSCs (Fig. 2A). In contrast, at the same FDR, we only classified 197 CpG loci (0.04% of the total sites tested) as DM between L-iPSCs and F-iPSCs. Even at substantially higher FDRs, the number of loci identified as DM between L-iPSCs and F-iPSCs was modest (Supplementary Tables 2A-D). Moreover, the 197 DM loci were not all independent; they clustered into 53 genomic regions, 37 of which are located near or within annotated genes (Fig. 2B).

**Figure 2.**
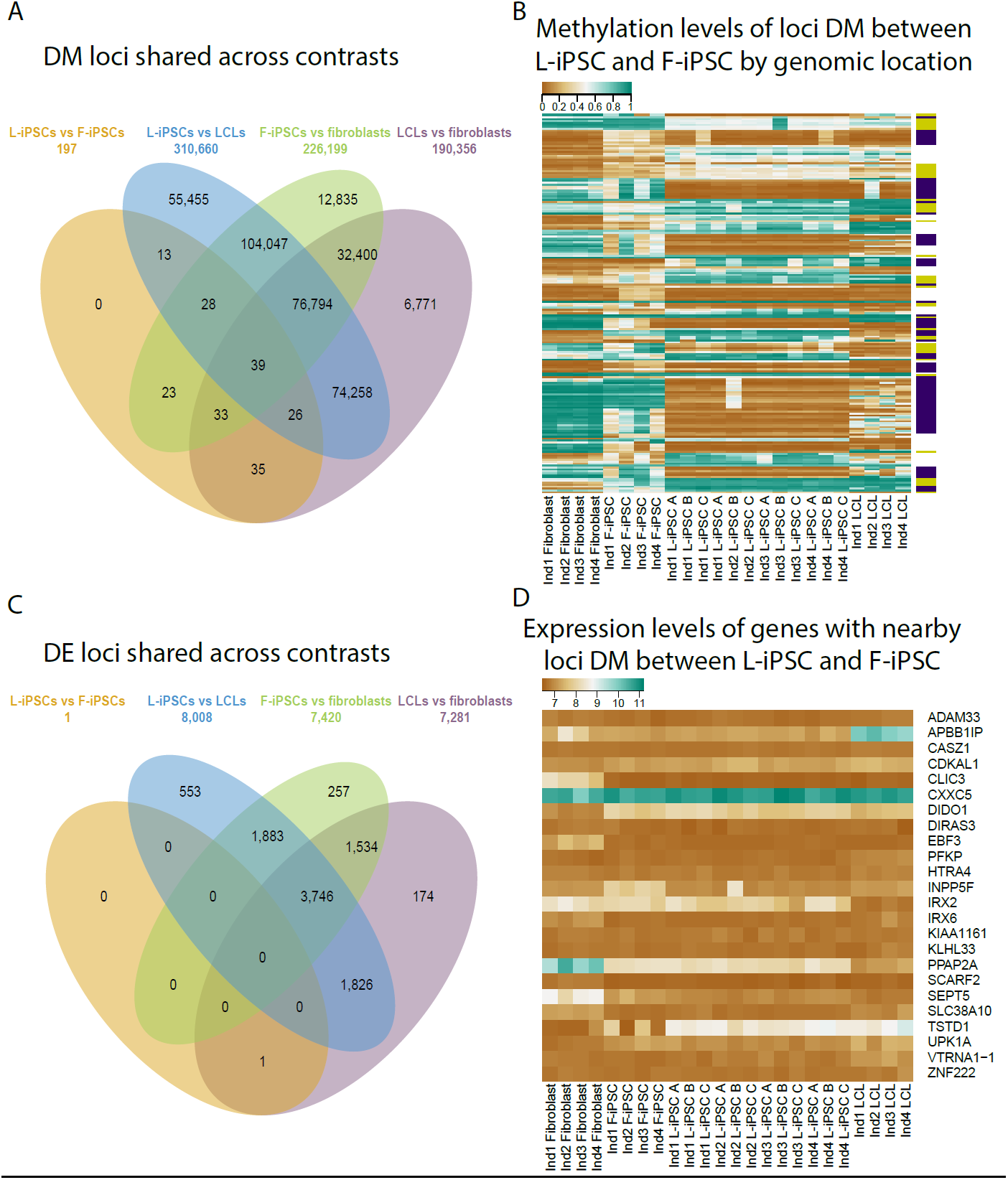
Differential methylation and gene expression between the four cell types (L-iPSC, F-iPSC, LCLs and fibroblasts). (a) A Venn diagram of differentially methylated (Dm) loci (FDR of 5%) overlapping between different contrasts. (b) A heat map of methylation levels at loci DM between L-iPSC and F-iPSC (n = 197), organized by genomic region (n = 53). The alternating purple and yellow color bars represent clusters of loci near a gene, while the white space represents loci in intergenic regions. (c) Venn diagram of differentially expressed (DE) genes (FDR of 5%) overlapping between different contrasts. (d) A heat map of expression levels for genes with nearby dM loci (where gene expression data were available; n= 24).

Of the 197 DM loci between L-iPSCs and F-iPSCs, 133 loci were also DM between LCLs and fibroblasts (a highly significant overlap; *χ*^2^ test; *P* < *10^-15^*). Moreover, 122 of these 133 DM loci showed a difference in methylation between LCLs and fibroblasts that was in the same direction as the one seen between L-iPSCs and F-iPSCs (sign test; *P* < *10^-15^*). In principle, these observations support the idea of epigenetic memory, namely that a subset of epigenetic differences between the somatic cells persists in the reprogrammed iPSCs. Yet our results indicate that epigenetic memory persists in a remarkably small number of loci.

We turned our attention to the gene expression data. We again used limma to identify (at an FDR of 5%) 7,281 differentially expressed (DE) genes between LCLs and fibroblasts, 8,008 DE genes between LCLs and L-iPSCs, and 7,420 DE genes between fibroblasts and F-iPSCs (Fig. 2C). In contrast, at the same FDR, we classified only a single gene (*TSTD1*) as DE between L-iPSCs and F-iPSCs. More generally, we found nearly no evidence for departure from a null model of no differences in gene expression levels between L-iPSCs and F-iPSCs (Supplementary Fig. 9A/B; Supplementary Tables 3A-D).

The single DE gene between L-iPSCs and F-iPSCs, *TSTD1* (*P = 6.28 × 10^-7^*; FDR 0.69%), is also DE between the LCLs and fibroblasts precursor cells. Moreover, 11 of 19 CpG sites that are located near the *TSTD1* gene, and are assayed by the methylation array, are among the 197 DM loci between L-iPSCs and F-iPSCs. We observed a decreased fold change of *TSTD1* expression when comparing between LCLs and fibroblasts (log2 fold change of 2.06) and L-iPSCs and F-iPSCs (log2 fold change of 1.34). This may be a case of epigenetic memory that maintains a gene expression residual difference, but it appears to be the only such case in our data. We found no evidence that any of the other DM loci are associated with gene expression differences between L-iPSCs and F-iPSCs (Fig. 2D). This is true even when we conservatively accounted for multiple tests by only considering the number of tests that involved genes that are associated with DM loci between L-iPSCs and F-iPSCs.

Our observations therefore indicate that remarkably little residual memory of the precursor somatic cell affects gene expression and methylation patterns in the reprogrammed iPSCs. To formally evaluate this we estimated the contribution of inter-individual differences and cell type of origin effects on variation in methylation and gene expression levels (see supplementary methods). The mean proportion of variance explained by donor individual is 11.2% and 9.3%, for the methylation and expression data while the mean proportion of variance explained by cell type of origin is -0.07% and 0.56% respectively (Fig. 3). The negative value is due to the adjustment of the estimated *R*^*2*^ to correct for variable degrees of freedom between covariates; it indicates that a model with cell type as a covariate fits the methylation data rather poorly (see Supplementary Fig. 11 for unadjusted and adjusted values).

**Figure 3.**
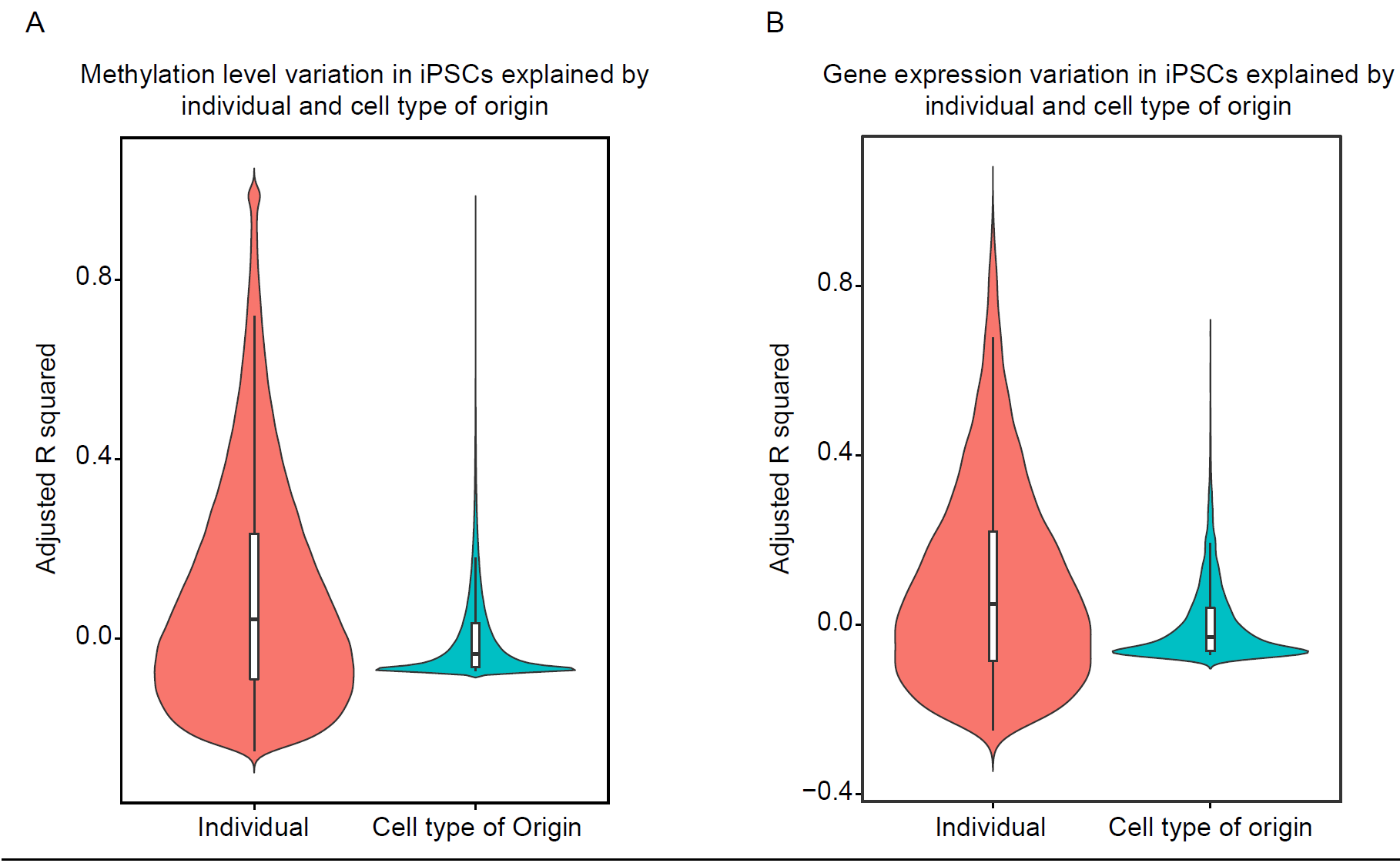
Estimated contribution of inter-individual differences and cell type of origin effects on variation in (a) methylation and (b) gene expression levels. Many individual loci have negative adjusted R^2^ values due to the applied correction for different numbers of parameters (see methods); negative values should be interpreted as an indication that the model fit the data poorly.

Put together, our analyses suggest that there is very little difference in DNA methylation and gene expression levels between L-iPSCs and F-iPSCs. In that sense, our results challenge the common view that epigenetic memory is prevalent in iPSCs. Yet, we believe that a careful examination suggests that our data are in fact consistent with previous studies. For example, though our interpretation is different because we have a benchmark of differences between individuals, we report a similar amount of DM between iPSCs derived from different cell types as seen in Kim et al. 2011^16^ Our study differs by explicitly modeling the contribution of genetic variation, whereas previous work has largely considered a single individual^12,14^ or confounded the effect of individual with cell type of origin^15,16^.

In conclusion, our study demonstrated that when accounting for individual, the impact of cell type of origin on DNA methylation and gene expression in iPSCs is limited to a small number of CpGs, which cluster into an even smaller number of genomic loci, and a single gene, with almost no detectable influence genome-wide. Our observations further confirm the usefulness of iPSCs for genetic studies regardless of the original somatic cell type. The high correlation of DNA methylation and gene expression levels (Supplementary Fig. 8C/D) between individuals (both within replicates and between L-iPSCs and F-iPSCs), demonstrate the faithfulness of the model. These data also suggest that future studies should focus on collecting more individuals rather than establishing multiple iPSC clones from the same individual.

## Methods

### Isolation and culture of fibroblasts and LCLs

Skin punch biopsies and blood were collected under University of Chicago IRB protocol 11-0524 (over three collection dates). All skin and blood samples per individual were processed in parallel. Fibroblast isolation and culture was conducted using the approach described in detail in Gallego Romero et al^25^. Briefly, skin punch biopsies (3mm) were digested using 0.5% collagenase B (Roche), isolated fibroblasts were cultured in DMEM (Life Technologies) supplemented with 10% fetal bovine serum (FBS; JR Scientific), 0.1mM NEAA, 2mM GlutaMAX (both from Life Technologies), 1% penicillin/streptomycin (Fisher), 64mg/L L-ascorbic acid 2-phosphate sesquimagnesium salt hydrate (Santa Cruz Biotechnology), at 5% CO_2_ and 5% O_2_.

All other cell culture was performed at 5% CO_2_ and atmospheric O_2_. For LCL generation, whole blood was drawn (within 20 minutes of obtaining skin punch biopsies) into two 8.5mL glass yellow top tubes (Acid Citrate Dextrose Solution A tubes; BD). Blood tubes were stored at room temperature and processed within 12 hours of collection. To isolate lymphocytes, we diluted whole blood with an equal amount of RPMI 1640 (Corning), diluted blood was slowly layered onto Ficoll-Paque (GE Lifescience) in 50 mL centrifuge tubes. This gradient was centrifuged at 1700 rpm for 30 minutes without acceleration or braking. Leukocytes and platelets formed a white band at the interface between the blood plasma and the Ficoll (called the buffy coat). We collected the buffy coat using a Pastette^®^ and to that added 10mL of PBS. The collected buffy coat was then washed three times with PBS.

For EBV transformation, 4 × 10^6^ fresh lymphocytes collected as described above were resuspended in a total of 4.5 ml of RPMI 1640 culture medium (Corning) containing 20% FBS and 1:100 phytohemagglutinin (PHA-M; LifeTechnologies) and transferred to a T-25 flask. EBV supernatant produced by the B95-8 cell lines (provided by the Ober lab) was added at 1:10 to the culture flask. Cells were left undisturbed for three to five days before adding fresh media. Flasks were subsequently examined weekly for changes in cell growth as indicated by acidic pH (yellow color) and the appearance of clumps of cells growing in suspension. Once growth was established (21-35 days), cells were diluted or split to several flasks. When the cell density reached 8 × 10^5^ to 1 × 10^6^ cells per mL they were cryopreserved at a density of 10 × 10^6^ cells per ml of freezing media in cryovials. All LCLs using this study were transformed with the same lot of EBV supernatant.

### Episomally-reprogrammed iPSCs

To establish iPSCs we transfected LCLs (Amaxa^™^ Nucleofector^™^ Technology; Lonza) and fibroblasts (Neon^®^ Transfection System; Life Technologies) with oriP/EBNA1 PCXLE based episomal plasmids that containing the genes *OCT3/4, SOX2, KLF4, L-MYC, LIN28*, and an shRNA against *p53*^19^. We supplemented these plasmids with an *in vitro*-transcribed EBNA1 mRNA transcript to promote exogenous vector retention following electroporation of the episomal vector^26, 27^ We plated a range of 10,000 - 40,000 transfected cells per well in a 6-well plate. Within 21 days colonies were visible and manually passaged onto a fresh plate of irradiated CF1 mouse embryonic fibroblasts (MEF). We passaged these new iPSC colonies on MEF in hESC media (DMEM/F12 (Corning) supplemented with 20% KOSR (LifeTechnologies), 0.1mM NEAA, 2mM GlutaMAX, 1% Pen/Strep, 0.1% 2-Mercaptoethanol (LifeTechnologies)). Fibroblast derived iPSCs were supplemented with 100ng/mL human basic fibroblast growth factor, versus 25ng/mL for LCL derived iPSCs; all other culture conditions were identical. After 10 passages of growth we transitioned the cultures to feeder-free conditions and cultured them for an additional three passages before collecting cell pellets for analysis. Feeder-free cultures were grown using 0.01 mg/cm^2^ (1:100) hESC-grade Matrigel (BD Sciences) and Essential 8 media (LifeTechnologies). Passaging was done using DPBS supplemented with 0.5mM EDTA. All RNA and DNA were isolated using Zymo dual extraction kits (Zymo Research) with a DNase treatment during RNA extraction (Qiagen).

### Characterization of iPSCs

All iPSC lines were characterized as described previously^25^. Briefly, we initially confirmed pluripotency using PluriTest^28^, a classifier that assigns samples a pluripotency score and novelty score based on genome-wide gene expression data. All samples were classified as pluripotent and had a low novelty score (Supplementary Fig. 1). We next performed qPCR using 1 μg of total RNA, converted to cDNA, from all samples to confirm the endogenous expression of pluripotency genes: *OCT3/4, NANOG,* and *SOX2* (Supplementary Fig. 2A-C). Additionally, we tested for the presence and expression of the EBV gene *EBNA-1* using PCR (primers and cycling conditions can be found in Supplementary Table 5) (Supplementary Fig. 2D/3). We tested all samples for both genomic integrations and vector-based EBV. We did this using primers designed to amplify the *EBNA-1* segment found in both the episomal vectors and the EBV used to transform LCLs. If the cell was positive (a single positive case was found: Ind4 F-iPSC), we further tested the origin of the EBV (genomic or episomal) using primers specific to the *LMP-2A* gene found in EBV or part of the sequence specific to the episomal plasmid (Supplementary Fig. 3). Finally, we confirmed the ability of all iPSC lines to differentiate into the three main germ layers using the embryoid body (EB) assay. The EBs were imaged for the presence of all three germ layers (Supplementary Fig. 4). In summary, all iPSC lines showed expression of pluripotent genes quantified by qPCR, generated EBs for all three germ layers, and were classified as pluripotent based on PluriTest.

### Processing of methylation array

Extracted DNA was bisulphite-converted and hybridized to the Infinium HumanMethylation450 BeadChip (Illumina) at the University of Chicago Functional Genomics facility. To validate the array probe specificity, probe sequences were mapped to an *in silico* bisulfite-converted genome using the Bismark aligner^29^. Only probes that mapped uniquely to the human genome were retained (n = 459,221). We further removed data from probes associated with low signal (detection *P-value > 0.01*) in more than 25% of samples (retained data from n = 455,910 loci). Raw output from the array (IDAT files) were processed using the minfi package^22^ in R.

We performed standard background correction as suggested by Illumina^22^, and corrected for the different distribution of the two probe types on the array using SWAN^23^ (Supplementary Fig. 5). Additionally, we quantile normalized the red and green color channels (corresponding to methylated and unmethylated signal respectively) separately (Supplementary Fig. 6A/B). To calculate methylation levels (reported as β-values) we divided the methylated signal by the total signal from both channels. β-values were considered estimates of the fraction of alleles methylated at that particular locus in the entire cell population.

### Processing of expression arrays

RNA quality was confirmed by quantifying sample’s RNA Integrity Number (RIN) on an Agilent 2100 Bioanalyzer (Agilent Technologies). All samples had a RIN of 10. The extracted RNA from all samples was hybridized to the Illumina HT12v4 Expression BeadChip array (Illumina) at the University of Chicago Functional Genomics facility. Sample processing was performed using the lumi package in R^30^. We excluded data from a subset of probes prior to our analysis: First, we mapped the probe sequences to the human genome hg19 and kept only those with a quality score of 37, indicative of unambiguous mapping (n = 40,198; note that we also explicitly prefiltered the 5,587 probes which were annotated as spanning exon-exon junctions to avoid mapping errors). Second, we downloaded the HapMap CEU SNPs (http://hapmap.ncbi.nlm.nih.gov/downloads/genotypes/2010-08_phaseI_I+III/forward/) and converted their coordinates from hg18 to hg19 using the UCSC liftOver utility^31^. We retained only those probes that did not overlap any SNP with a minor allele frequency greater than 5% (n = 34,508). Third, we converted the Illumina probe IDs to Ensembl gene IDs using the R/Bioconductor package biomaRt^32^ and retained only those probes that are associated with exactly one Ensembl gene ID (Ensembl 75 - Feb 2014; n = 22,032). The full pipeline was implemented using the Python package Snakemake^33^. We defined a gene as expressed in a given sample if at least one probe mapping to it had a detection *P-value* < 0.05. In the case of L-iPSCs, we defined a gene as expressed in an individual if any associated probes had a detection *P-value* < 0.05 in at least one biological replicate. Using these criteria, we identified all genes expressed in at least three individuals in at least one cell type (Supplementary Fig. 7; n = 14,111 probes associated with 11,054 annotated genes). In the case that multiple expressed probes were associated with the same ENSEMBL gene (n = 3,057), we only retained data from the 3’-most detected probe. Following these filtration steps, we obtained estimates of expression levels in all samples across 11,054 genes. Data from the 11,054 genes were quantile normalized using the lumiExpresso function in lumi^30^ (Supplementary Fig. 6C/D).

### Unsupervised hierarchical clustering and heatmaps

Data from only autosomal probes were retained for the hierarchical clustering analyses in order to reduce bias towards clustering by individual or sex (n = 10,648 expression, and n = 445,277 methylation). We calculated a matrix of pairwise Euclidean distances between samples from the methylation and expression data separately. From these matrices we performed hierarchical clustering analyzing using the complete linkage method as implemented in the R function hclust. The 1,000 most variable loci were defined by taking the loci with the highest variance in iPSCs. Clustering based on the 1,000 most variable probes were processed in an identical manner as above. Heatmaps were generated from matrices of pairwise Pearson correlations between samples.

### Analysis of differences in gene expression and methylation levels

Data from probes on both autosomes and sex chromosomes were included in this analysis, given that individuals were balanced across cell types (n = 455,910 CpGs; n = 11,054 genes). Additionally, we anticipated that sites on the sex chromosomes may be particularly sensitive to mis-regulation during reprogramming^34^. Differential expression and methylation analyses were performed using linear modeling and empirical Bayes methods as implemented in the limma package^24^. We tested for differential methylation and expression, using locus-specific models, between L-iPSCs and F-iPSCs; L-iPSCs and LCLs; F-iPSCs and fibroblasts; and between fibroblasts and LCLs. We considered a locus DM or DE at an FDR < 5% (Benjamini Hochberg). We also tested for DE genes between L-iPSCs and F-iPSCs using only genes that were classified as DE between L-iPSCs and LCLs; F-iPSCs and fibroblasts; and LCLs and fibroblasts (Supplementary Fig. 9A). We estimated FDRs separately each time we considered only subsets of the data.

Due to the imbalance of L-iPSC samples to F-iPSC samples we repeated our analyses using data from a reduced set of samples. Namely, we randomly sampled a single replicate of the L-iPSC from each individual. As expected, reducing the number of L-iPSC samples greatly reduces the number of loci classified as DM between L-iPSCs and F-iPSCs as well as between L-iPSCs and LCLs. However, the number of DM loci was reduced across all other contrasts as limma models the entire matrix together (Supplementary Fig. 10). Interestingly, we found that different combinations of replicates yielded DE genes other than *TSTD1*. Therefore, we sampled all possible combinations and overall, found six genes that were classified as DE (FDR 5%; see Supplementary Table 4) in at least one of the combinations of reduced samples. Of note, we never classify *TSTD1* as DE (FDR 5%) in the reduced data set. The most common DE gene, *INPP5F*, is the only gene that also has nearby DM CpGs (five of the 25 nearby loci). Additionally, in the full model, *INPP5F* has the second lowest *P value* (uncorrected *P = 6.84 × 10^-5^*; FDR 38%). However, *INPP5F* was not DE between LCLs and fibroblasts, but was DE between LCLs and L-iPSCs and also fibroblasts and F-iPSCs (Supplementary Tables 3A-D; Fig. 2D).

### Enrichment of DM loci in regulatory and genomic features

We employed two strategies to identify enrichments of DM loci between L-iPSCs and F-iPSCs in regulatory features. First, we used the regulatory states defined by Ernst et al.^35^. We tested for enrichments in all regulatory categories using a *χ*-square test comparing the number DM loci and total probes within each regulatory class to the number DM loci and total probes outside the regulatory class. We found no significant enrichment for any of the defined regulatory states.

Next, we used the UCSC_RefGene_Group annotation as supplied by Illumina. These annotations detail the location of probes in relation to genes (1st Exon, 3’ UTR, 5’ UTR, Gene Body, within 1.5kb of a TSS or within 200bp of a TSS). We identified significant enrichments of DM loci within 1.5kb of a TSS and gene bodies. However, there are six probes classified as both within a gene body and within 1.5kb of a TSS. We chose to report both results because it is difficult to deconvolute these categories.

We also considered the position of DM loci in relation to genes. The annotations were defined by Illumina. We were able to identify 37 genes associated with DM loci, but we only had corresponding gene expression data for 24 of these genes. We attempted to identify signals of enrichment in DE levels between L-iPSCs and F-iPSCs in these 24 genes. To this end, we compared the log fold changes in gene expression between L-iPSCs and F-iPSCs from genes with nearby DM loci between L-iPSCs and F-iPSCs to 10,000 random samplings of log fold change in expression between L-iPSCs and F-iPSCs from all genes and found no enrichment for increased log fold changes.

### Proportion of variance explained

To estimate the proportion of variance explained by individual and cell type of origin we performed an ANOVA using the effect estimates from a multiple linear regression containing fixed effects for individuals and cell type of origin. Only data from autosomes were included in this analysis so that the results would not be biased toward differences in individuals (n = 10,648 expression, and n = 445,277 methylation). For each term we calculated the proportion of variance explained by dividing the sum of squares for that term by the total sum of squares for all terms. However, given that our two parameters had differing degrees of freedom (n = 3 for the individual and n = 1 for cell type of origin) we were concerned that the proportion of variance explained by individual was artificially inflated. To account for this, we performed the following correction to the R^2^ from the individual and cell type of origin terms:

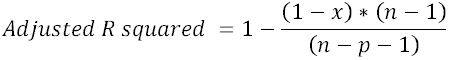

where n is the number of individuals and p is the number of parameters being estimated (Supplementary Fig. 11)^36^. This correction yields many individual loci with negative R^2^ values (see Fig. 3), but results in a more unbiased estimate of the mean across all loci.

## Acknowledgments

We would like to thank John D. Blischak as well as members of the Pritchard, Stephens, and Gilad laboratories for valuable discussions during the preparation of this manuscript. This work was supported by Howard Hughes Medical Institute funds to J.K.P. and by NIH grants MH084703 and HG006123 to Y.G. and J.K.P. C.L.K. was supported by a CTSA TL1 pre-doctoral fellowship. N.E.B. was supported by an NIH training grant GM007197 and an NIH pre-doctoral award F31 AG 044948. I.G.R. was supported by a Sir Henry Wellcome Postdoctoral Fellowship. The expression and methylation data are available at GEO under accession GSE65079.

## Contributions

C.L.K., B.J.P., J.K.P., and Y.G. developed the concept and designed experiments. I.G.R. developed the IRB for primary cell collection and oversaw cell collection. B.J.P. cultured all primary tissues and made F-iPSCs. C.L.K. and K.P. made L-iPSCs and did all iPSC QC. C.L.K. analyzed expression data and N.E.B analyzed methylation data. N.E.B., C.L.K., and Y.G. wrote the manuscript with input from all authors.

